# Electrospray sample injection for single-particle imaging with X-ray lasers

**DOI:** 10.1101/453456

**Authors:** Johan Bielecki, Max F. Hantke, Benedikt J. Daurer, Hemanth K. N. Reddy, Dirk Hasse, Daniel S. D. Larsson, Laura H. Gunn, Martin Svenda, Anna Munke, Jonas A. Sellberg, Leonie Flueckiger, Alberto Pietrini, Carl Nettelblad, Ida Lundholm, Gunilla Carlsson, Kenta Okamoto, Nicusor Timneanu, Daniel Westphal, Olena Kulyk, Akifumi Higashiura, Gijs van der Schot, Duane Loh, Taylor E. Wysong, Christoph Bostedt, Tais Gorkhover, Bianca Iwan, M. Marvin Seibert, Timur Osipov, Peter Walter, Philip Hart, Maximilian Bucher, Anatoli Ulmer, Dipanwita Ray, Gabriella Carini, Ken R. Ferguson, Inger Andersson, Jakob Andreasson, Janos Hajdu, Filipe R. N. C. Maia

## Abstract

The possibility of imaging single proteins constitutes an exciting challenge for X-ray lasers. Despite encouraging results on large particles, imaging small particles has proven to be difficult for two reasons: not quite high enough pulse intensity from currently available X-ray lasers and, as we demonstrate here, contamination of the aerosolised molecules by non-volatile contaminants in the solution. The amount of contamination on the sample depends on the initial droplet-size during aerosolisation. Here we show that with our electrospray injector we can decrease the size of aerosol droplets and demonstrate virtually contaminant-free sample delivery of organelles, small virions, and proteins. The results presented here, together with the increased performance of next generation X-ray lasers, constitute an important stepping stone towards the ultimate goal of protein structure determination from imaging at room temperature and high temporal resolution.

## Introduction

Coherent diffractive imaging^1^ with femtosecond ultrabright pulses from X-ray free-electron lasers (XFELs)^2,3^ has been successfully applied to large viruses, organelles, and even entire cells^4,5,6^. Critical for the success of these pioneering studies were high-fluence XFEL beams, specialised detectors, low background noise, and efficient sample delivery into the XFEL focus^4,5,7^. The most widely used injector for this approach, the Uppsala injector^8^, generates a droplet aerosol by atomising the sample solution with a gas-dynamic virtual nozzle (GDVN)^9^. The volatile droplet components evaporate, leaving behind one aerosol particle for every occupied droplet. A skimmer removes excess aerosol carrier gas and an aerodynamic lens focuses the aerosol to a narrow beam that is directed into the XFEL focus for imaging individual aerosol particles. While this injector has been used for imaging biological particles with diameters between 80 nanometers^10^ and 2000 nanometers^6^, imaging smaller particles has proven challenging^11,12^. Particles appeared rounder, larger, and showed a higher level of polydispersity than in solution^11,12,5^

It has been suspected that large and polydisperse initial droplets may be the cause for this size and shape mismatch^12^. Non-volatile contaminants are often unavoidable components of the sample solution and the initial droplet size determines how much remains attached to the aerosolised particle after solvent evaporation. This problem is also known in electrospray-ionisation mass spectrometry as these contaminants degrade the mass spectral signal-to-noise ratio^13^.

For droplet formation with GDVNs, a narrow cone-jet from the nozzle of a capillary is hydrodynamically tapered by a He sheath gas, up to the point at which the jet becomes unstable and breaks up into small droplets. This jet-atomisation technique is efficient for the continuous creation of a large number of aerosol droplets with diameters of micrometers to sub-micrometers^9^.

Electrospray (ES) is an alternative jet-atomisation technique^14,15^. By applying a voltage to the liquid the jet is squeezed into a Taylor cone, by electrostatic forces without the requirement of exerting pressure by a sheath gas. ES has become a very powerful method to aerosolise biological particles with a wide range of sizes for examination by mass spectrometry^16,17^ or differential mobility analysis (DMA)^18^. Low flow rates and small droplets can be obtained, achieving gentle aerosolisation with low contamination. A prerequisite for a stable Taylor cone is an inert and dielectric ambient gas that does not react and does not remove electrical charge from the liquid. A mixture of CO_2_ and N_2_ at a pressure of at least 800 mbar fulfills this requirement^16^, which in our injector leads to a mass flow of 1.2 standard litre per minute (SLM) N_2_ and 0.15 SLM CO_2_. In contrast, the GDVN produces less than 0.5 SLM of He.

## Results

We modified the design of the Uppsala aerosol sample injector and substituted the GDVN with an ES aerosoliser. To reduce the increased mass flow from the dielectric gas we added an additional nozzle-skimmer stage (Fig. 1a). The operational parameters for the GDVN aerosoliser are significantly different from the ES aerosoliser (Table 1). While our GDVNs are operated at liquid flow rates (*Q*) on the order of µL/min and our ES aerosoliser is operated at ~20 times lower flow rates. As the droplet volume (*V*) of the ES aerosoliser is ~300 times smaller than for GDVNs, ES produces droplets at ~15 times higher rate (*R*=*Q*/*V*). Theoretically, therefore, higher hit rates should be achievable by ES compared to GDVN aerosolisation under usual conditions.

**Table 1.**
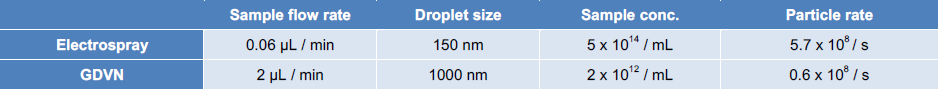
Aerosolisation parameters. Characteristic parameters for sample aerosolisation with electrospray and a gas-dynamic virtual nozzle (GDVN) assuming an average droplet occupancy of 1.

**Figure 1.**
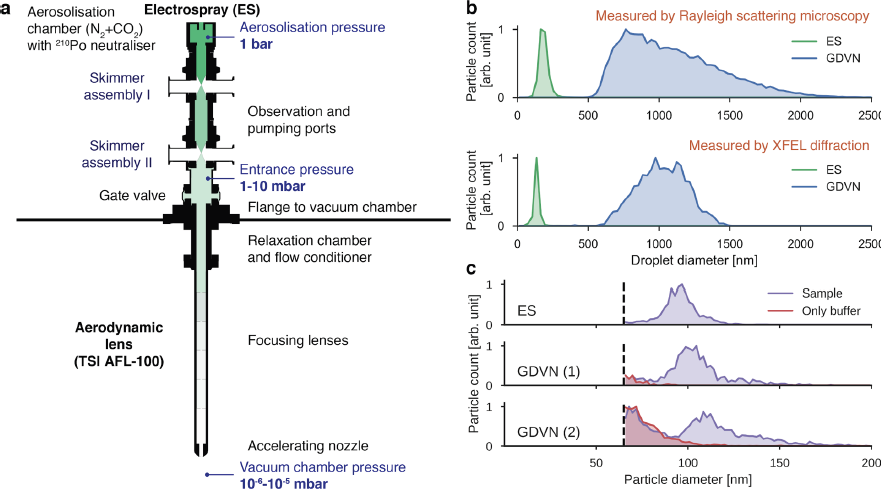
Electrospray aerosol injector. **a** Design of the electrospray (ES) aerosol injector. In the aerosolisation chamber the ES nebuliser generates droplets that are neutralised with a ^210^Po alpha emitter. The ES nebuliser is operated in an atmosphere of N_2_ and CO_2_ at 1 bar. The aerosol is transported through two nozzle-skimmer assemblies where excess gas is pumped away. At a reduced pressure of 1-10 mbar the aerosol enters the aerosol lens-stack, which focuses it to a narrow particle beam entering the experimental chamber, which is held at 10^-6^-10^-5^ mbar pressure to match requirements for XFEL imaging. **b** Size distributions of initial droplets for ES (green) and GDVN (blue) aerosols determined by RSM (top) and by XFEL diffraction (bottom). The results of the two sizing methods are comparable within the limits of reproducibility expected for the manually manufactured nozzles and variations in operational parameters, such as pressures, voltage, and flow rate. **c** RSM size distributions of aerosolised particles from carboxysome sample (purple) and from its buffer solution (red). Data collected on electrosprayed particles are shown in the first panel (95 nm median, 14 nm FWHM) and data collected on particles injected by GDVN at two different pressure configurations (Table 2) are shown in the second (102 nm median, 17 nm FWHM) and third panel (105 nm median, 17 nm FWHM). Dashed lines indicate the detection limit.

To compare droplet formation between ES and GDVN, we first determined the size distributions of initial droplets of the two aerosolisers (Fig. 1b) by measuring the size of particles that are formed when injecting sucrose solution^19^. Sizes were measured by Rayleigh scattering microscopy (RSM)^8^ and, in addition, by XFEL diffraction^5,12^. The droplets generated with the GDVN span a wide range of diameters (500 to 2000 nm) whereas droplets generated with the ES aerosoliser are smaller, and more monodisperse (100 to 200 nm).

For comparing bioparticle aerosols generated with the two injector designs, we selected carboxysomes as a biological test sample. Carboxysomes are polyhedral cell organelles that are heterogeneous in size with an average diameter of about 100 nanometers^5^. Using RSM we found that particles have, on average, larger diameters if aerosolised with a GDVN compared to ES (purple histograms, Fig. 1c). This observation confirms that the amount of non-volatile contaminants that accumulates on the surface of aerosol particles increases with the size of the initial droplet. Furthermore, control measurements on only buffer (red histograms in Fig. 1c) revealed the presence of contaminant particles in the GDVN aerosols. These are likely aggregates of non-volatile buffer remaining after solvent evaporation from empty droplets.

We tested the ES injector for X-ray imaging at the Atomic, Molecular & Optical science (AMO) beamline at the Linac Coherent Light Source (LCLS). As test samples, we selected carboxysomes, tomato bushy stunt virus (TBSV) particles, and the protein ribulose-1,5-bisphosphate carboxylase/oxygenase (Rubisco, EC 4.1.1.39).

In a previous study with GDVN aerosolisation we obtained high-quality diffraction images on carboxysomes, albeit most of the particles appeared round instead of icosahedral as would be expected^5^. From the new diffraction data with ES aerosolisation (5,000 hits recorded within 7 minutes), we reconstructed projection images of carboxysomes (Fig. 2a) and determined the size distribution (Fig. 2b). Almost all particles matched projections of an icosahedral particle and both the median and standard deviation of the size distribution are in agreement with our RSM measurements (Fig. 1c). These results confirm that ES injection, in comparison to GDVN aerosolisation, reduces the amount of non-volatile contaminants.

**Figure 2.**
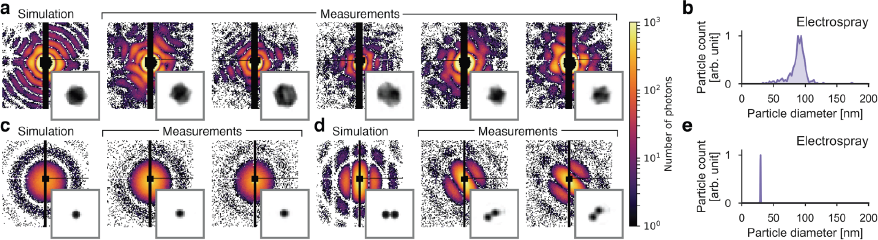
XFEL diffraction data of biological particles injected with the electrospray aerosol injector. **a** Simulated and measured diffraction patterns of carboxysomes and **b** their size distribution (90 nm median, 13 nm FWHM) determined from the measured diffraction patterns. **c, d** Simulated and measured diffraction patterns of TBSV particles (**c** singles, **d** clusters of two) and **e** their size distribution (30 nm median, 1 nm FWHM) determined from the measured diffraction patterns. Insets in panels **a**, **c** and **d** show 2D projection images reconstructed from the respective diffraction patterns. The edge length of the insets corresponds to 220 nm.

TBSV particles are monodisperse with a diameter of about 35 nm. Despite their small size, 6,000 high-quality diffraction patterns of single and double particles (Fig. 2c,d) were collected within one hour of data collection. Particle clusters are expected due to the high sample concentration and the possibility of double-occupancy of the droplets. The reconstructed projection images show the expected shape. The size distribution has a full-width at half-maximum (FWHM) smaller than 1 nm (Fig. 2e), which shows that ES did not alter the size distribution of the sample.

As a third test sample, we injected 11-nm sized Rubisco proteins. The X-ray cross section for a Rubisco protein is about 30 times smaller than for a TBSV particle. In Fig. 3a we compare the predicted signal (red dashed lines), using the measured incident peak intensity, to radially averaged diffraction data on injected Rubisco and respective control data on injected sample buffer solution, injection gas, and a dark run (solid lines in panel 1, 2, 3, and 4, respectively). The comparison shows that the predicted diffraction pattern for a single protein was too faint to exceed gas background fluctuations. Nevertheless, we found diffraction patterns that exceeded the amplitude of background fluctuations and two examples are shown in Fig. 3b. From the diffraction images, we determined particle diameters matching the approximate size of a protein cluster of 2 to 3 particles (Fig. 3c).

**Figure 3.**
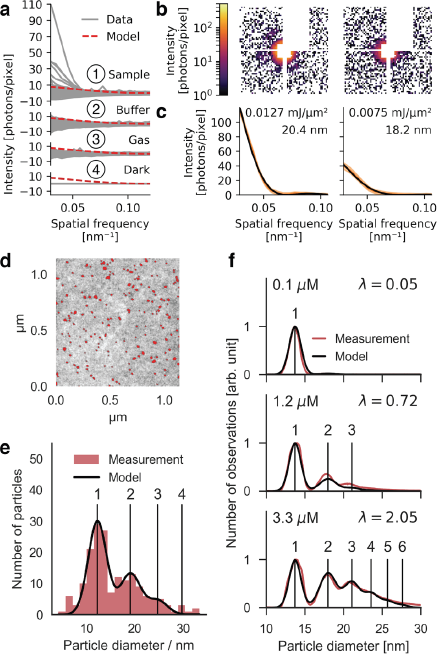
Injection of Rubisco proteins. **a** Radial averages of background subtracted diffraction patterns recorded during injection of sample (1), buffer solution (2), only gas (3), and during a dark run (4), respectively. **b** Diffraction patterns of two intense sample hits. **c** Radial averages (orange lines) of the diffraction patterns shown in **b** and fits (black lines) to a sphere model that best match the data. Light orange areas indicate the confidence intervals of the data (+/- one standard deviation). The fit values for intensity and sphere diameter are annotated. **d** STEM image of Rubisco proteins injected onto a TEM sample support film. Detected particles are highlighted in red. **e** The red histogram shows the distribution of particle diameters derived from **d**. The black line shows the fit of our droplet occupancy model to the data. The good match indicates that the electrosprayed proteins were successfully transferred into the interaction region. **f** DMA data of electrosprayed Rubisco proteins at three concentrations. Our droplet occupancy model (black) was fitted to the measured size histograms (red). The agreement shows that by changing concentration we specifically control the protein cluster composition.

Due to the weak scattering signal obtained with the pulse intensity that was available at the LCLS we could not conclusively determine if single Rubisco proteins were delivered into the interaction region. To answer this question, we injected Rubisco and deposited the injected particles for examination by scanning transmission electron microscopy (STEM) (Fig. 3d). We extracted the size distribution of deposited particles (red histogram in Fig. 3e) by integrating the areas of particles in the image. The size distribution matched a Poissonian droplet occupancy model (black line in Fig. 3e), which proves that we injected single Rubisco proteins into vacuum. We confirmed the validity of this model by measuring the size distribution of the same sample at a range of concentrations by DMA (Fig. 3f).

In conclusion, we report successful single particle imaging of 35 nm biological samples - significantly smaller than previously possible. Our adaptation of the Uppsala injector for ES was shown to decrease droplet sizes and was shown to enable delivery of single proteins into vacuum. With this achievement we overcome one of the major experimental hurdles that has hindered progress for XFEL imaging of small biological particles. For large particles, the smaller droplets of ES are also beneficial as they reduce contamination from non-volatile buffer components. As a result of the higher reproducibility of aerosolised particles, ES injection is also expected to increase attainable resolution in 3D reconstructions.

Further development in lens stack design^20^ and aerosolisation geometry is expected to increase the particle transmission and decrease fluorescence and scattering background from injection gas that dominated the noise in our diffraction data. We anticipate that such diffraction data from single proteins will be possible to analyse using established 3D reconstruction methods^21,10^. The results presented here together with the increased X-ray flux and repetition rate of next generation FEL facilities such as European XFEL and LCLS II, will constitute an important stepping stone towards the ultimate goal of protein structure determination from imaging at room temperature and high temporal resolution.

## Methods

### Sample preparation

#### Sucrose solutions for aerosol droplet size determination

Initial droplet size distributions of ES and GDVN aerosols were determined by measuring the size distributions of particles generated by injecting sucrose solution. The particle diameter d_p_ is related to the initial droplet diameter d_0_ via the relation d_p_=d_0_c^1/3^, where c is the v/v concentration of sucrose. Sucrose concentrations were adjusted to achieve final particle sizes after solvent evaporation of around 100 nm, suitable for sizing by RSM and XFEL diffraction. For the RSM measurements we used sucrose solutions at concentrations of 12 % (v/v) for ES and 0.1 % (v/v) for GDVN aerosolisation and for the XFEL diffraction measurements 5 % (v/v) for ES and 0.1 % (v/v) for GDVN injection.

#### Carboxysome purification

Carboxysomes were purified from *Halothiobacillus neapolitanus* DMS15147 cells as previously described in ref. 5 with minor changes to the protocol with respect to the lysis of the cells (omitting the sonication step). After harvesting by centrifugation, the cells were resuspended in 50 ml TEMB-lysozyme buffer (10 mM Tris-HCl pH 8.0, 10 mM MgCl_2_, 20 mM NaHCO_3_, 1 mM EDTA, 100 μg/mL lysozyme, pH 8.0). The cell suspension was mixed with 50 mL of B-PER^®^ Bacterial Protein Extraction reagent (Thermo SCIENTIFIC) and incubated for about 10 minutes at room temperature on a rotary shaker. When the solution turned viscous, due to DNA release from broken cells, deoxyribonuclease I from bovine pancreas (SIGMA-Aldrich) was added to a final concentration of 1 μg/ml. The suspension was incubated for an additional 30 minutes at room temperature on a rotary shaker. After pelleting the debris, the carboxysomes were purified by centrifugation and resuspension as described in ref. 5. For ES injection we used the purified sample at a concentration of about 10^13^ particles per mL in TEMB-lysozyme buffer (i.e. without exchanging the buffer). For GDVN injection carboxysomes were buffer exchanged by eluting the sample into 20 mM ammonium acetate solution (pH 7.5) using a PD MiniTrap G-25 column (GE Healthcare). This exchange was performed twice. We followed the same buffer exchange protocol for the control measurements on TEMB-lysozyme buffer.

#### Tomato bushy stunt virus purification

Tomato bushy stunt virus (strain BSV-3, American Tissue Culture Collection code PV-90) was propagated in *Nicotiana benthamiana* grown at 25 °C under a 16/8-hour light/dark cycle. Leaves were mechanically inoculated using carborundum and virus extract. At 6–8 days post infection, leaves that showed severe signs of infection were harvested and stored at −20°C. Frozen leaves, chilled with liquid nitrogen, were ground into a fine powder using a mortar and pestle and transferred into an ice-cooled Bead-beater (BioSpec Products Inc; 2-mm zircona beads). Ice-cold extraction buffer (50 mM sodium phosphate, pH 5, 1 mM TCEP) was added to the ground leaf tissue in a ratio of 5:1 (v/w), before five rounds of 60/60 second on/off cycles. The solution was cleared from precipitated proteins and cell debris by centrifugation at 8,000 *g* for 30 minutes at 4 °C. The supernatant was sequentially filtered using 5 µm, 0.8 µm and 0.2 µm syringe filters. Virus particles were sedimented by ultracentrifugation at 100,000 *g* for 2 hours at 4 °C. The resulting pellets were carefully resuspended into native buffer (50 mM Tris-HCl, pH 7.5, 20 mM CaCl_2_) and cleared from any undissolved particulates by centrifugation for one minute at 20,000 *g*. The re-suspended pellet was floated on a 15–60% pre-formed sucrose gradient (made using native buffer) and was subjected to rate-zonal centrifugation at 100,000 *g* for 2 hours at 4 °C. The virus particles could be seen as a band approximately 1/3 from the top of the tube when illuminated from the top. The band was recovered in fractions by pipetting and analyzed for UV absorption at 260 and 280 nm (NanoDrop; ThermoFisher Scientific) and by SDS-PAGE. The sucrose was removed by dialysis into native buffer. Exchange into the injection buffer (25 mM ammonium acetate, pH 5) was achieved by multiple rounds of sample dilution and subsequent concentration using a VivaSpin 10,000 MWCO concentrator (Vivascience). The final particle concentration used for injection was 3–5 x 10^14^ mL^-1^. Sample quality was verified by measuring size homogeneity and shape by DLS (W130i; AvidNano, Ltd) and negative-stain electron microscopy (FEI Quanta; ThermoFisher Scientific).

#### Rubisco purification

*Spinacia oleracea* Rubisco was purified as previously described in ref. 22. After long-term storage at −80°C, the sample was further purified by size-exclusion chromatography using a Hiload 26/60 Superdex 200 (GE healthcare) column attached to a NGC chromatography system (BioRad). Separation was performed at 4 °C, with a flow rate of 2 mL/min, in Superdex buffer (50 mM Tris-Cl, pH 8.0, 100 mM NaCl, 1 mM EDTA). Peak fractions containing Rubisco identified by SDS-PAGE (data not shown) were pooled, and concentrated using a VivaSpin 30,000 MWCO concentrator (Vivascience).

Purified *S. oleracea* Rubisco was incubated at room temperature for 30 min in the presence of equimolar 4-carboxy-D-arabinitol-1,5-bisphosphate (4-CABP), a reaction-intermediate analogue that binds tightly and irreversibly to Rubisco active sites. 4-CABP binding induces a conformational change of a surface exposed loop to cover the active site of Rubisco, thereby reducing the structural heterogeneity of the sample^22^.

Prior to injection, the protein was buffer exchanged into Ammonium Acetate Sample Buffer (20 mM ammonium acetate, pH 7.97) over a PD10 desalting column (GE Healthcare) as described above.

### Sample aerosolisation

#### Gas-dynamic virtual nozzles (GDVNs)

GDVNs were manufactured in-house according to the general design presented in ref. 9. The generation of sub-micrometer droplets require a large reduction in gas pressure around the liquid jet meniscus together with a low liquid flow rate. To achieve this, we utilised a ‘flush’ geometry as described in ref. 24 together with a 20 µm inner diameter liquid capillary, whose tip was conically grinded at approximately 15-20° attack angle. Stable jets were achieved with liquid flow rates between 0.5-2 µL/min and outer He sheath flow between 0.5-1.5 SLM.

#### Electrospray (ES)

The ES nebuliser was based on the design introduced in ref. 25. The sample was supplied with 360 µm outer diameter fused silica capillaries with inner diameter of 30 µm when injecting TBSV, sucrose and Rubisco, while capillaries with an inner diameter of 40 µm were used when injecting carboxysomes. The capillaries were conically grinded at 30° attack angle until the tip of the capillary had a ‘plateau’ with a diameter of 80 µm. During nebulization, the tip of the capillary was positioned approximately 1 mm away from a grounded orifice plate with 0.5 mm orifice diameter. The formation of a Taylor cone was achieved by applying a 2-3 kV voltage to the sample inside the sample reservoir while the sample was flowing with a 50-100 nL/min flow rate. The flow rate was achieved by applying 1-10 psi of overpressure in the sample reservoir. To keep the Taylor cone stable, an influx of 0.15 L/min CO_2_ + 1 L/min N_2_ was necessary in order to avoid de-charging of the liquid at the meniscus. The exact voltages and flow rates needed to achieve a stable Taylor cone vary with the conductivity of the sample. In this configuration, stable operation could be achieved with conductivities between 1700-7000 µS/cm. The charged droplets generated by the ES aerosolisation were neutralised with a Po-210 alpha source.

### Aerosol injection

Particles were delivered into the in-vacuum interaction region for GDVN aerosolisation with the original and for ES aerosolisation with the modified version of the Uppsala aerosol injector^4,5,26^. Excess gas from the aerosolisation process was removed in a nozzle-skimmer stage located between the aerosolisation compartment and the aerodynamic lens stack. For GDVN aerosolisation^9,27^, a single nozzle-skimmer stage, with 0.3 mm and 0.6 mm skimmer aperture, was required in order to reduce the gas load inside the aerodynamic lens stack. To accommodate the increased mass flow for ES aerosolisation we added a second nozzle-skimmer stage (skimmer assembly I in Fig. 1a), with 0.8 mm nozzle and 1 mm skimmer apertures. This additional stage was located upstream of the existing stage (skimmer assembly II in Fig. 1a). In both stages, the nozzle-skimmer distance was set such that the skimmer was located within the zone of silence^26^ of the freely expanding gas exiting the nozzle.

### Particle sizing

#### Particle sizing by differential mobility analysis (DMA)

DMA measurements were carried out with the TSI3080 electrostatic classifier together with the TSI3081 differential mobility analyzer. The ES aerosol described above was used as input to the electrostatic classifier, while the size-selected particle output was detected with the TSI3786 condensed particle counter. In all, this system enabled detection and relative concentration measurements of particles 10-1000 nm in diameter.

#### Particle sizing by Rayleigh scattering microscopy (RSM)

RSM data were acquired as described in ref. 8. Size calibration was carried out with suspensions of Monodisperse Polystyrene Sphere Size Standards (Fischer Scientific, NIST traceable size standard, refractive index 1.5983). The calibration factors were rescaled on the basis of estimates for the refractive index of the respective particle species (carboxysomes: 1.4 [ref. 28], sucrose 1.5376 [ref. 29]).

#### Particle size determination from XFEL diffraction intensities

The sizes of injected sucrose, carboxysome and TBSV particles were determined by fitting the diffraction image of a uniform sphere model to the measured diffraction patterns^12^. Table 2 lists the data sets that were used. Prior to fitting, the diffraction patterns were truncated below 0.5 photons and pixels were binned (sucrose and TBSV data 6 by 6, carboxysome data 4 by 4). Throughout the fitting procedure a binary mask was used that excluded hot, saturated and shadowed pixels, and pixels at large diffraction angles where the signal from non-spherical objects is expected to deviate significantly from the sphere model. All run-specific parameters can be found in the files *amol3116_sizing.csv* and *amol3416_sizing.csv* under the open repository *https://github.com/mhantke/electrospray_injection*. In a last refinement step, we modified the fitting model to include an offset term to account for uniform background that was observed in the diffraction data. The sizing was carried out in an automated fashion together with a manual inspection of the fitted results and discarding of failed fits.

**Table 2.**
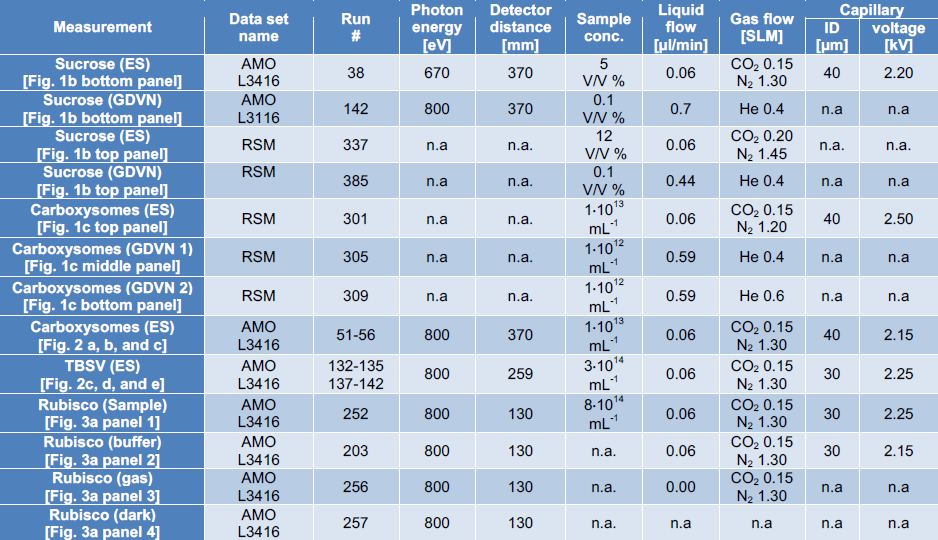
Data sets used for this study.

Rubisco particles were sized by fitting the radial diffraction intensities of a sphere model to the radially averaged diffraction intensities of the measurement. To validate our results, we checked that the incident intensity that resulted from the fit fell into the range of intensities expected for the X-ray beam focus^30^. For this calculation we assumed that the particles had the, for proteins characteristic, mass density of 1.35 g/cm^3^ and the atomic composition of H_86_C_52_N_13_O_15_S (ref. 3).

#### Particle sizing by electron microscopy

Rubisco particles exiting in a collimated beam from the aerosol injector were collected by streaking on a 400 mesh Cu F/C EM Grid (Ted Pella, Inc). The grid was then imaged without staining at 240k magnification in a FEI Quanta FEG 650 using a STEM detector at an acquisition time of 1 µs and at an acceleration voltage of 30 kV.

### XFEL diffraction measurements

#### Data collection

XFEL diffraction data were collected inside the LAMP chamber^31^ at the AMO endstation^32^ of the LCLS. The particle beam exiting from the Uppsala aerosol injector was intersected with the X-ray beam. The LCLS generated X-ray pulses of 1-2 mJ at 800 eV photon energy (1.55 nm wavelength) with a pulse duration of 170 fs and a peak fluence of 0.02 mJ/µm^2^ (ref. 14) at a repetition rate of 120 pulses per second. About 5% of the LCLS pulses were dumped (“BYKICK” mode) in order to monitor continuously the dark background. This means that the LCLS delivered effectively only about 114 pulses per second to the interaction region. Diffraction images were recorded synchronously with a pair of pnCCD area detector panels^33^ operated in gain mode 5. The panels were placed at distances of 250 mm (TBSV data) and 370 mm (carboxysome and sucrose data). Each panel has a sensitive area of 76.8 mm x 38.4 mm with 1024 x 512 pixels. The direct beam and small-angle scattering passed through the gap between the panels. At 250 mm detector distance the gap was 3.3 mm wide and at 370 mm detector distance it was 5.5 mm wide. Data were monitored live with the Hummingbird software package^34^

#### Data pre-processing

Diffraction data were pre-processed using the Hummingbird software package^34^ and Psana^35^. Configuration files (*conf_preproc.py* and *conf_amol3416.py*) can be downloaded from https://github.com/mhantke/electrospray_injection. The data sets that were used for analysis are listed in **Supplementary Table 2**. First, raw data were pedestal subtracted using dark frames and rescaled to the unit of X-ray photons. Pedestal correction was following by a 3-step common mode subtraction procedure that was carried out for each panel individually, first for every quadrant (half panel), then for each fast, and finally for each slowly changing pixel dimension. Common mode is defined as the median pixel value of the selection of pixels that measure below 0.5 photons. For the faulty top-right quadrant additionally ASIC-wise common mode subtractions were applied, first for the fast then for the slowly changing pixel dimension. For certain runs (defined in *amol3116_run_params.csv* and *amol3416_run_params.csv*) all pixels of the inner one or two ASICs of the faulty quadrant were upscaled by a factor of two. Detector geometry was applied by taking into account the relative position of the detector halves, and pnCCD read-out timing issue for particular runs, and the column mismatch that was caused by a wiring error of the pnCCD chip. As hits, we selected those diffraction patterns which counted more than 3500 pixels measuring at least one photon and being located further than 200 pixels away from the center.

#### Data prediction

Diffraction data for carboxysomes, TBSV particles, and Rubisco proteins were simulated with the Condor software package^36^. For Rubisco proteins the electron density was estimated to 0.43 Å^-3^ on the basis of 1.35 g/cm^3^ mass density and H_86_C_52_N_13_O_15_S atomic composition for proteins^2^. The incident intensity was set to the measured peak fluence of 0.02 mJ/µm^2^ (ref. 30).

#### Image reconstruction

For retrieving the phase of selected carboxysome and TBSV diffraction patterns and reconstructing 2D projection images, we used the *Hawk* software package^37^. Prior to phasing, the diffraction patterns were truncated at 0.5 photons and binned to 128×128 images. We used a binary mask excluding hot, saturated and shadowed pixels.

The support was initialised with a static spherical mask of radius slightly larger than the expected particle size. The iterative phase retrieval was performed with 1000 iterations of the Relaxed averaged alternating reflections (RAAR) algorithm^38^ (TBSV hits) or Hybrid input output (HIO) algorithm^39^ (carboxysome hits) algorithm followed by 1000 iterations of the error reduction (ER) algorithm^39^ in both cases enforcing the projected electron densities to be real and positive. The final reconstruction is an average of 100 independent reconstructions with a random initial guess for the phases. To check for reproducibility of the reconstructions we calculated phase retrieval transfer functions (PRTFs) (Fig. 4).

**Figure 4.**
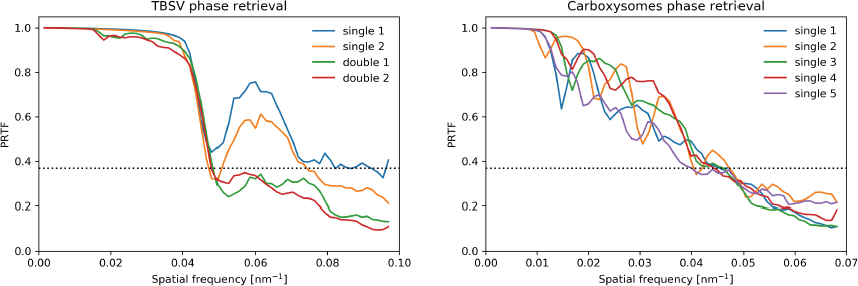
Phase retrieval transfer functions (PRTFs) for reconstructed projection images shown in Fig. 2a, c, d. The dashed lines indicate the value *e*^-1^, often used as threshold for judging the reproducibly of the retrieved phases.

#### Rubisco data analysis

Diffraction patterns were pre-processed as described above and then binned 16 by 16 pixels to improve the signal-to-noise ratio. Then pixel values below the background floor of half a photon were set to zero to reduce the background from gas fluorescence and visible light and all other values were rounded to the closest integer value. For every pixel the variance and the mean value were calculated from the buffer run. Pixels for which the ratio of variance and mean value deviated by less than 0.3 from 1 were identified as good pixels because of the indication that their values followed Poisson statistics. Pixels that did not fall into this category were masked out. The mask was extended manually to exclude the halo of the direct beam and the edges of the detector quadrants. Finally, images were background corrected by subtraction of the median read-out value for every pixel, respectively.

### Droplet occupancy model

The droplet occupancy by particles during droplet formation was modelled as a Poissonian process. The expectation value *λ* for the occupancy *n* of a droplet is given by the product of particle concentration in solution and droplet volume. For multiply occupied droplets (*n*>1) the particles stick together and form a (non-specific) complex. We assumed that the diameter of the complex *d_n_* does not grow as *d_n_*~n^1/3^ because the new complex will be most likely less compact than a sphere. Instead we fitted the distributions shown in Fig. 3e and 3f by using the scaling law *d_n_*~n^1/a^ with the free parameter *a<3*. We obtained *a*=1.57 for the deposited proteins imaged by STEM and *a*=2.56 for the DMA data.

## Acknowledgements

This work was supported by the Swedish Research Council, the Knut and Alice Wallenberg Foundation, the European Research Council, the Röntgen-Ångström Cluster, the projects Advanced research using high intensity laser produced photons and particles (ADONIS) (CZ.02.1.01/0.0/0.0/16_019/0000789) and Structural dynamics of biomolecular systems (ELIBIO) (CZ.02.1.01/0.0/0.0/15_003/ 0000447) from the European Regional Development Fund, the Swedish Foundation for Strategic Research, the Swedish Foundation for International Cooperation in Research and Higher Education (STINT), and the Wellcome Trust (204732/Z/16/Z).

J.A acknowledges support from the Ministry of Education, Youth and Sports as part of targeted support from the National Programme of Sustainability II and the Chalmers Area of Advance; Materials Science. The contribution of O.K. is part of the EUCALL project funded from the EU Horizon 2020 research and innovation programme under grant agreement No 654220.

C.B. and M.B. were supported by the U.S. Department of Energy, Office of Basic Energy Sciences, Division of Chemical Sciences, Geosciences, and Biosciences under contract DE-AC02-06CH11357. B.I. and T.G. are grateful for the support from the Volkswagen Foundation through the Peter-Paul Ewald fellowship.

Use of the Linac Coherent Light Source (LCLS), SLAC National Accelerator Laboratory, is supported by the U.S. Department of Energy, Office of Science, Office of Basic Energy Sciences under Contract No. DE-AC02-76SF00515.

## Author Contributions

J.B., M.F.H., B.J.D., J.A., J.H. and F.R.N.C.M. conceived and designed the experiments. J.B., M.F.H., and D.W. designed the adaptation of the Uppsala injector for electrospray. H.K.N.R., D.H., D.S.D.L, L.H.G., M.S., A.M., A.H., and I.A. prepared samples. D.S.D.L. and O.K. manufactured nozzles for sample aerosolisation. J.B., M.F.H., B.J.D., H.K.N.R, D.H., D.S.D.L, L.H.G., M.S., A.M., J.A.S., L.F., A.P., C.N., I.L., G.Carlsson, K.O., N.T., D.W., O.K., A.H., G.v.d.S., D.L., T.E.W., C.B., T.G., B.I., M.M.S., T.O., P.W., P.H., M.B., A.U., D.R., G.Carini, K.R.F., I.A., J.A., J.H., and F.R.N.C.M carried out the experiments. J.B., M.F.H. and B.J.D. analysed the data with input from F.R.N.C.M. J.B., M.F.H., B.J.D., and F.R.N.C.M. prepared the manuscript with input from all other coauthors.

